# The Mia40 substrate Mix17 exposes its N-terminus to the cytosolic side of the mitochondrial outer membrane

**DOI:** 10.1101/2024.05.30.596364

**Authors:** Moritz Resch, Johanna S. Frickel, Korbinian Dischinger, Rachel Choo Shen Wen, Kai Hell, Max E. Harner

## Abstract

Mitochondrial architecture and the contacts between the outer and the inner mitochondrial membrane depend on the mitochondrial contact site and cristae organizing system (MICOS) that is highly conserved from yeast to human. Mutations in the mammalian MICOS subunit Mic14/CHCHD10 have been linked to amyotrophic lateral sclerosis and frontotemporal dementia, indicating the importance of this protein. Mic14/CHCHD10 has a yeast ortholog, Mix17, a protein of unknown function, which has not been shown to interact with MICOS so far. As a first step to elucidate the function of Mix17 and its orthologs, we analyzed its interactions, biogenesis and mitochondrial sublocation.

We report that Mix17 is no stable MICOS subunit in yeast. Our data suggest that Mix17 is the first Mia40 substrate in the mitochondrial outer membrane. Unlike all other Mia40 substrates, Mix17 spans the outer membrane and exposes its N-terminus to the cytosol. The insertion of Mix17 is likely to be mediated by its interaction with Tom40, the pore of the TOM complex. Moreover, we show that the exposure of Mix17 to the cytosolic side of the membrane depends on its N-terminus.

## Introduction

The mitochondrial contact site and cristae organizing system (MICOS) is essential for the organization of mitochondrial architecture (John et al. 2005, Rabl et al. 2009, Harner et al. 2011, Hoppins et al. 2011, von der Malsburg et al. 2011). Moreover, it forms contacts between the inner and the outer membrane. The importance of the MICOS complex is reflected by its conservation from yeast to mammalians (Alkhaja et al. 2012). The mammalian subunits have six homologs in the yeast *Saccharomyces cerevisiae:* Mic10, Mic12, Mic19, Mic26, Mic27 and Mic60 (Harner et al. 2011, Hoppins et al. 2011, von der Malsburg et al. 2011). Five of the MICOS components are integral inner membrane proteins facing into the IMS and follow the presequence import pathway (Harner et al. 2011, Hoppins et al. 2011, von der Malsburg et al. 2011, Ueda et al. 2019). Only the Mic19 component of MICOS, an IMS protein that is peripherally associated with the inner membrane, employs the Mia40-dependent import pathway (Ueda et al. 2019).

The mammalian MICOS complex contains additional subunits. Recently, the CHCHD10 protein, also termed Mic14, was shown to interact with Mic60/mitofilin, Mic19 and Mic25 (Genin et al. 2016). Interestingly, CHCHD10/Mic14 is highly conserved. It has an ortholog in yeast, Mix17, whose association with MICOS has not been analyzed so far. Also, its function remains unclear. The deletion of Mix17 negatively affects mitochondrial respiration by an unknown mechanism (Longen et al. 2009). Similar to Mic14, Mix17 contains a CHCH domain with the twin Cx_9_C motif of typical Mia40 substrates and its import was shown to be Mia40-dependent (Gabriel et al. 2007, Longen et al. 2009, Gornicka et al. 2014). In contrast to typical Mia40 substrates with twin Cx_9_C motif, Mix17 contains additional conserved sequence features, which may play a crucial role for its biogenesis pathway.

Here, we analyzed protein interactions of Mix17, its submitochondrial location and the import pathway of Mix17 as an important step to elucidate the functional role of Mix17 in mitochondria. In contrast to the reported interaction of CHCHD10/Mic14 and MICOS (Genin et al. 2016), yeast Mix17 is not stably associated with MICOS. Interestingly, we demonstrate that Mix17 is accessible to proteases added to intact mitochondria. Unlike all other Mia40 substrates, Mix17 apparently inserts into the outer membrane, suggesting that it is a novel kind of Mia40 substrate. We show that Mix17 spans the outer membrane, most likely through the TOM complex. Moreover, the N-terminus of Mix17 is essential for the insertion of Mix17 into the outer membrane and, in addition, improves the import of Mix17 into mitochondria.

## Results

### Mix17 is no subunit of the MICOS complex in yeast

The interaction of the potential mammalian MICOS subunit ChChD10/Mic14 is still a matter of debate (Genin et al. 2016, Burstein et al. 2018), which might be explained by the suggested cell-type-specific function of the protein (Burstein et al. 2018). Since this protein is highly conserved from yeast to human (Fig. S1) (Longen et al. 2009), we first addressed whether Mix17 interacts with the MICOS complex in the unicellular organism *Saccharomyces cerevisiae*.

To this end, we applied immunoprecipitation using mitochondria of yeast strains expressing 3xHA tagged versions of Mic10 or Mic60. With both tagged proteins, we were able to co-isolate known MICOS subunits, indicating that the complex remained stable under these conditions. Mix17, however, did not co-precipitate with Mic10 or Mic60 (Fig. 1A). Next, we tested whether it is possible, to co-isolate MICOS subunits via Mix17. Therefore, we purified Mix17 using an antibody specific to the endogenous protein. Although we could efficiently immunoprecipitate Mix17, the MICOS subunits Mic10, Mic26, Mic27 and Mic60 were not co-isolated (Fig. 1B).

**Fig. 1.**
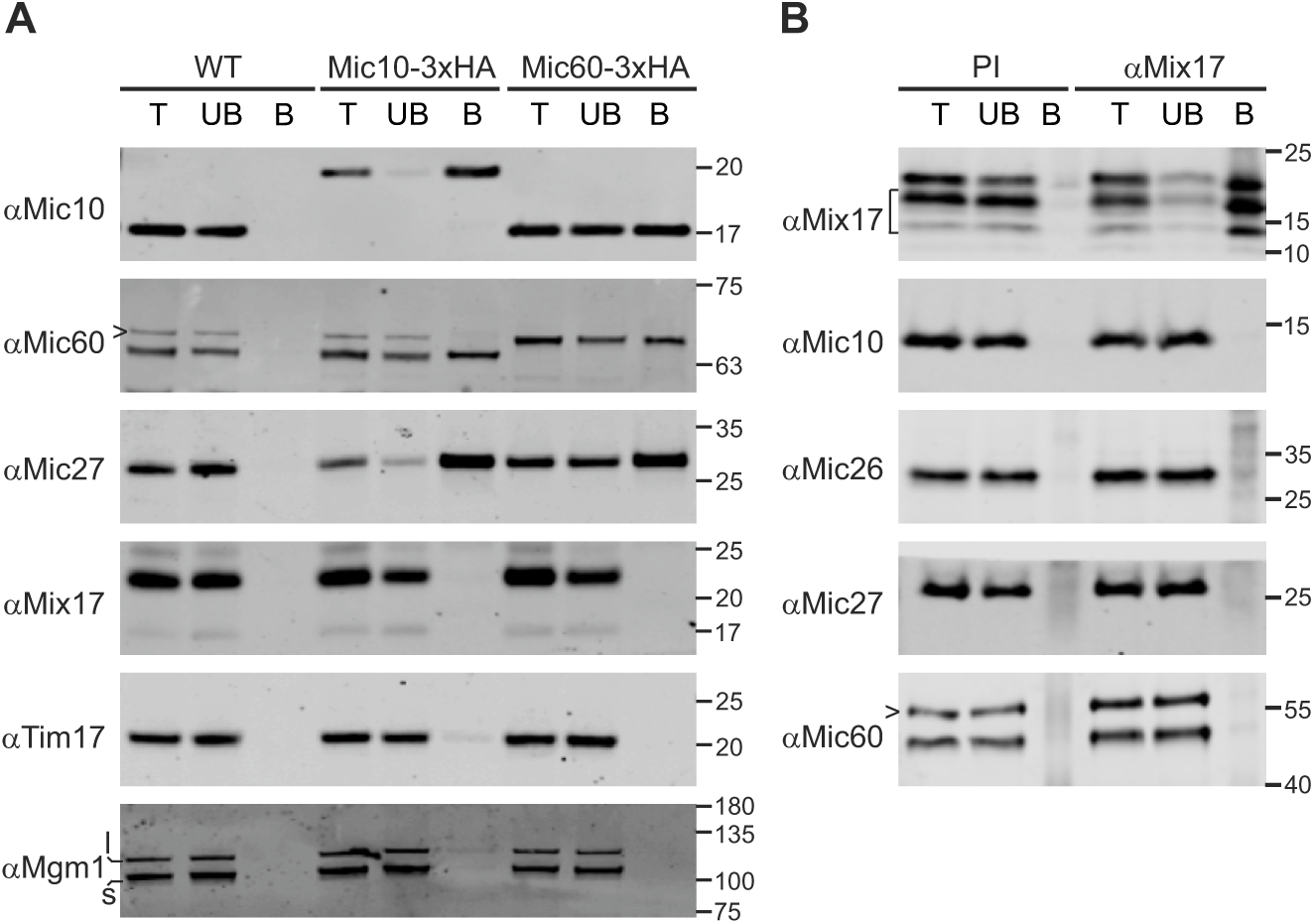
Mix17 is no MICOS subunit. **(A)** Mitochondria of wild type yeast or yeast strains expressing Mic10-3xHA or Mic603xHA were lysed in digitonin containing buffer (1% w/v). Lysates were subjected to immunoprecipitation using anti-HA affinity agarose. Samples were analyzed by SDS-PAGE and immunoblotting. T, total lysate (4%); UB, unbound protein (4%); B, bound protein (100%). l, long isoform of Mgm1; s, short isoform of Mgm1. Arrowhead, cross reaction of the anti-Mic60 antibody. **(B)** Wild type mitochondria were treated as in (A) and subjected to immunoprecipitation using a Mix17 specific antibody or pre-immune serum bound to protein Asepharose. Samples were analyzed by SDS-PAGE and immunoblotting. T, total lysate (4%); UB, unbound protein (4%); B, bound protein (100%). PI, pre-immune serum; bracket, degradation products of Mix17; arrowhead, cross reaction of the anti-Mic60 antibody. Immunoblots are representative of three repeats.

Taken together, we conclude that Mix17, at least under the conditions tested, does not stably associate with MICOS in yeast.

### Mix17 is exposed to the cytosol with a N_out_-C_in_ topology

Next, we analyzed the location of Mix17 in isolated mitochondria. Surprisingly, we found that Mix17, like the outer membrane marker Tom70, is easily degradable by proteinase K (PK) in intact mitochondria (Fig. 2A). Addition of PK resulted in the generation of two smaller fragments, which still could be recognized by an antibody against the C-terminal Myc-tag. This indicates that Mix17 spans the outer membrane exposing its N-terminus to the cytosol (Fig. 2A). As control for intact mitochondria, the inner membrane protein Tim50 was stable at these conditions. When we treated osmotically swollen mitochondria (SW) with PK, Mix17 and Tim50 were completely gone (Fig. 2A). The strong signal reduction of Mix17 upon re-isolation of swollen mitochondria as well as its presence in the soluble fraction (S) upon alkaline extraction suggests that Mix17 spans the outer membrane, possibly in a proteinaceous environment, rather than being integrated into the lipid bilayer of the membrane (Fig. 2A). As expected, the integral membrane proteins Tim50 and Tom70 stayed in the membrane protein fraction (M) upon alkaline extraction and were not lost upon swelling and re-isolation (Fig. 2A). Importantly, we obtained virtually the same results for the endogenous Mix17 protein in wild type mitochondria indicating that the Myc-tag did not alter the topology (Fig. 2B). Next, we confirmed our results with another protease. Mix17 contains 6 arginine residues in its N-terminus (Suppl. Fig. S1), which should be recognized by trypsin, if the N-terminus is exposed to the cytosol. Indeed, we detected a faster migrating fragment upon trypsin treatment of intact mitochondria. At the same time, Tom70 was degraded but Tim50 stayed intact (Fig. 2C). Next, we asked whether we could reproduce this stunning result following the *in vitro* import of radioactively labeled Mix17 precursor protein. Incubation of Mix17 precursor with energized wild type mitochondria allowed its efficient import into isolated mitochondria. Moreover, addition of trypsin to intact mitochondria after the import reaction resulted in the generation of the same fragment as detected for the endogenous protein (Fig. 2D).

**Fig. 2.**
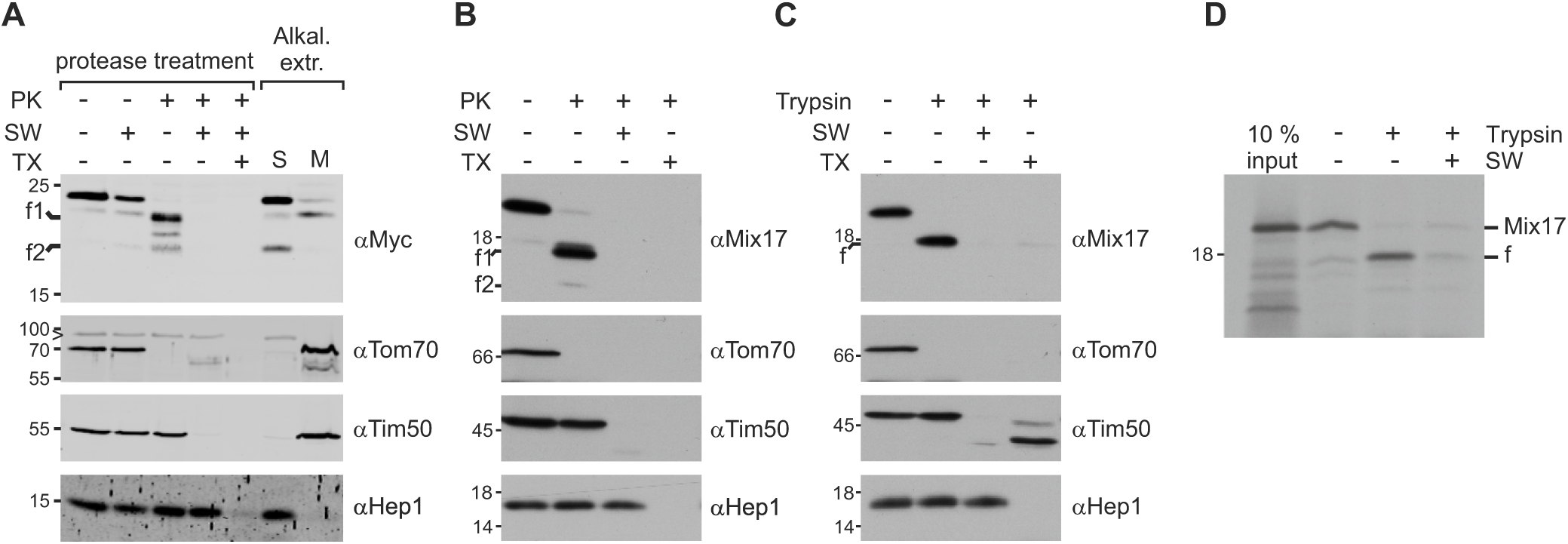
Mix17 has the ability to span the outer membrane with an N_out_-C_in_ topology. **(A)** Mix17 exposes its N-terminus to the cytosol. Mitochondria isolated from a Mix17-3xMyc expressing strain were subjected to protease and alkaline treatments. (Left) Mitochondria were left at isotonic conditions, osmotically swollen (SW), or lysed in Triton X-100 containing buffer (TX). Proteinase K (PK) was added as indicated. f1, f2, fragments generated by PK. (Right) Mitochondria were treated with alkaline buffer to separate soluble (S) and membrane proteins (M). Samples were analyzed by SDS-PAGE and immunoblotting. Arrowhead, cross reaction of the anti-Tom70 antibody. **(B)** Wild type Mix17 shows the same topology as Mix17-3xMyc. Yeast wild type mitochondria were treated and subjected to PK treatment as described in (A). **(C)** The N-terminus of Mix17 is sensitive to trypsin. Wild type mitochondria were subjected to protease treatment using trypsin. Samples were analyzed as in (A). f, fragment generated by trypsin. Blots are representative of at least three repeats. **(D)** *In vitro* imported Mix17 adopts the same topology as the endogenous protein. Mix17 protein was synthesized *in vitro* in the presence of ^35^S-methionine and incubated with isolated mitochondria. Mitochondria were re-isolated and treated as in (C). The samples were analyzed by SDS-PAGE and autoradiography. The autoradiograph is representative of at least three repeats. The full-length protein (Mix17) or the fragment generated by trypsin (f) are indicated. 10% input, 10% of the amount of radiolabeled protein added to each lane.

In summary, our results showed that Mix17 spans the mitochondrial outer membrane in a N_out_-C_in_ topology. Moreover, the loose association of Mix17 to the outer membrane indicates that it might be embedded in the outer membrane in a proteinaceous environment.

### Mia40 and the twin CX9C motif of Mix17 are essential for the import of Mix17

Next, we set out to analyze which factors are important for the import of Mix17 and its insertion into the outer membrane. In line with previous results (Gabriel et al. 2007, Gornicka et al. 2014), we show that Mia40 is essential for the import of Mix17 despite its uncommon mitochondrial sublocation for a Mia40 substrate. Import of Mix17 was strongly impaired in mitochondria depleted of Mia40 (Fig. 3A). The import of Mia40 substrates depends on a conserved CX9C motif, which is also present in Mix17 and its homologs (Suppl. Fig. S1). A disulfide bond is formed between these cysteines by the MIA40 system, which is essential for the import of this class of proteins (Mesecke et al. 2005, Hell 2008, Koehler and Tienson 2009, Riemer et al. 2009, Stojanovski et al. 2012, Edwards et al. 2020). In accordance with this, inhibition of the formation of a disulfide bond by dithiothreitol (DTT) strongly reduced the import efficiency of Mix17 into wild type mitochondria (Fig. 3B). The introduction of point mutations into Mix17 had an even more obvious effect. The import of precursor protein, in which the cysteines of the twin CX9C motif were replaced by serines, was almost completely inhibited (Fig. 3C). Finally, we tested whether Mia40 is also essential for the import of Mix17 *in vivo*. We isolated mitochondria from wild type cells and Mia40-depleted cells and tested for the steady state level of Mix17 in comparison to control proteins and wild type mitochondria. Interestingly, Mix17 was virtually absent upon downregulation of Mia40, similar to the Mia40 substrate Tim13. In contrast, the level of mitochondrial proteins that do not depend on Mia40 did not change significantly (Fig. 3D).

**Fig. 3.**
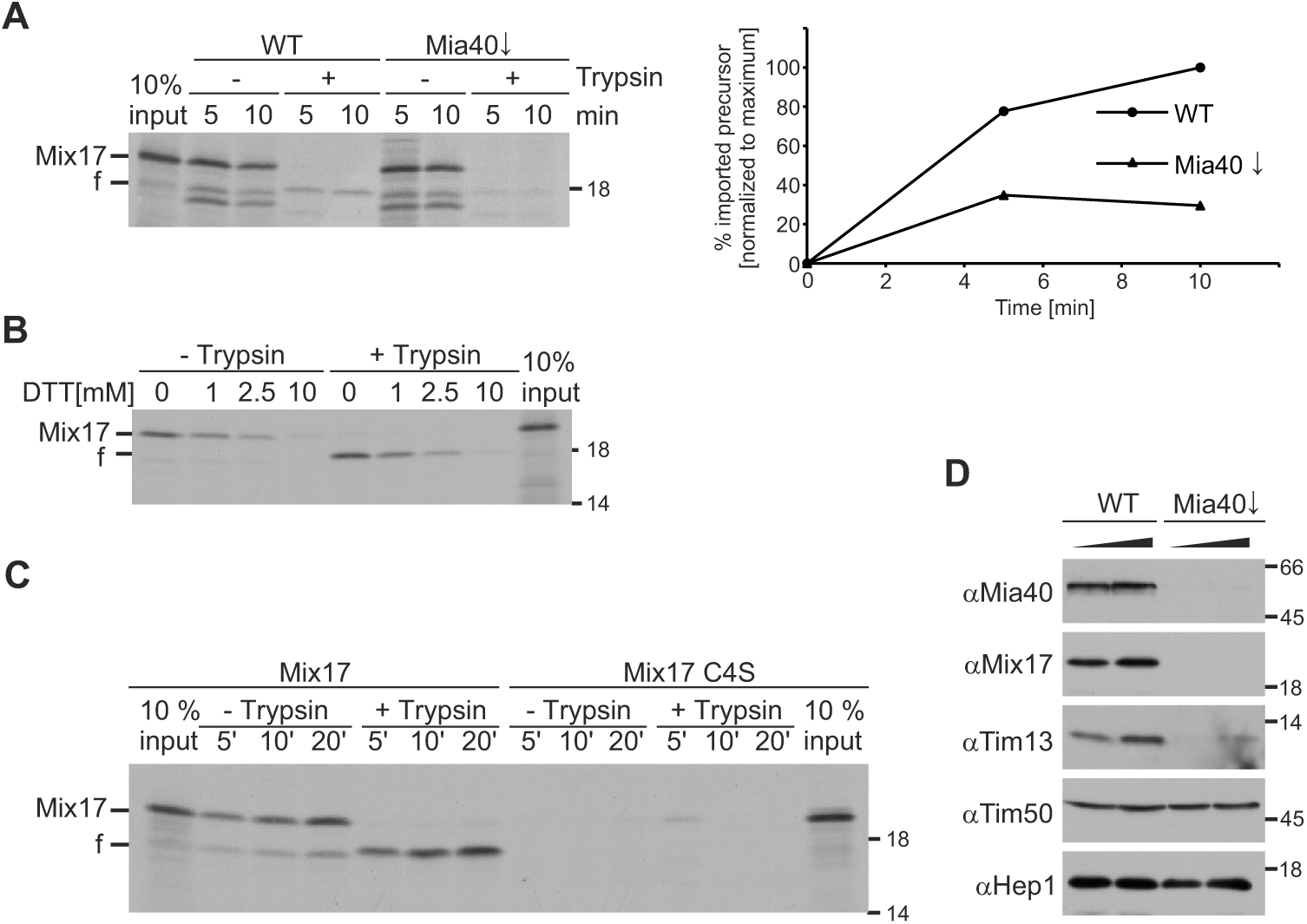
Mia40 is essential for the import of Mix17. **(A)** The import of Mix17 depends on Mia40. ^35^S-labeled Mix17 protein was *in vitro* translated and incubated with mitochondria isolated from wild type yeast or a mutant strain, in which Mia40 was downregulated (Mia40↓). Samples were taken at the indicated time points and treated with trypsin as indicated. Mitochondria were re-isolated and the samples were analyzed by SDS-PAGE and autoradiography. f, fragment generated by trypsin. The autoradiograph of the experiment (left) and the quantification are shown (right). **(B)** DTT inhibits the import of Mix17. ^35^S-labeled Mix17 protein was imported into wild type mitochondria for 20 min in presence of increasing concentrations of DTT. Samples were analyzed as in (A). **(C)** The conserved CX_9_C motif is essential for the import of Mix17. ^35^S-labeled Mix17 protein and a mutant lacking the four highly conserved C-terminal cysteine residues (Mix17 C4S) were incubated with wild type mitochondria. Samples were treated with trypsin and analyzed as in (A). **(D)** The steady state level of endogenous Mix17 depend on Mia40. Wild type and Mia40↓ mitochondria were isolated and 25 and 50 µg of mitochondrial proteins were analyzed by SDS-PAGE and immunoblotting using the indicated antibodies. Blots and autoradiographs are representative of at least two experiments.

Taken together, in line with previous results (Gabriel et al. 2007, Gornicka et al. 2014), we demonstrate that the disulfide relay system and the twin CX9C motif of Mix17 are essential for its import into mitochondria.

### The N-terminus of Mix17 is essential for the insertion into the outer membrane and improves the import into mitochondria

Mix 17 contains, in addition to the twin Cx_9_C-motif, further characteristic sequence features. Its N-terminus has a predicted probability to be a mitochondrial targeting signal (Claros and Vincens 1996). Therefore, we asked whether the N-terminus plays a crucial role for the import of Mix17 and its insertion into the mitochondrial outer membrane. Presequence proteins harboring a mitochondrial targeting signal depend on mitochondrial membrane potential for their import. Thus, we first analyzed whether the import of Mix17 is membrane-potential-dependent.

We observed similar import efficiencies in presence and absence of membrane potential (Fig. 4A). The integration of the N-terminus of Mix17 into the outer membrane was also not affected in absence of membrane potential. Imported Mix17 was still accessible to trypsin. Next, we asked, whether the N-terminal segment per se is important for the import of Mix17. Therefore, we deleted the first 24 amino acids of Mix17 and tested its import efficiency compared to full-length Mix17. The truncated Mix17 Δ1-24 variant showed a reduced import efficiency to 20% of the one of the full-length protein (Fig. 4B). Of note, the radiolabeled Mix17 Δ1-24 lysate was considerably stronger than that of the full-length version. When we treated mitochondria with trypsin after the import reaction, we were not able to detect a reduction in the size of Mix17 Δ1-24 (Fig. 4B). This was expected, since the truncation led to the loss of the trypsin cleavage site that was required for detection of the trypsin accessibility. To be able to analyze the insertion of Mix17 Δ1-24 into the outer membrane, we repeated the experiment using PK instead of trypsin. Again, we detected an import defect of the truncated version. Similar to previous experiments (Fig. 2A), treatment of mitochondria with PK generated two fragments of imported full-length Mix17. Such PK fragments were not observed following the import of Mix17 Δ1-24. Importantly, both PK-fragments migrated considerably faster upon SDS-PAGE than Mix17 Δ1-24. If Mix17 Δ1-24 inserted into the outer membrane, this variant, therefore, should have been accessible to the added PK (Fig. 4C). This strongly indicates that the N-terminus of Mix17 is essential for insertion of Mix17 into the outer membrane. Consistently, we did not detect clipping of Mix17 Δ1-24 by PK or trypsin, when we tested mitochondria isolated from a *MIX17 Δ1-24-FLAG* expressing yeast strain in the protease accessibility assays (Fig. 4D). Thus, the N-terminus appears to be also crucial *in vivo* for the insertion of Mix17 into the outer membrane.

**Fig. 4.**
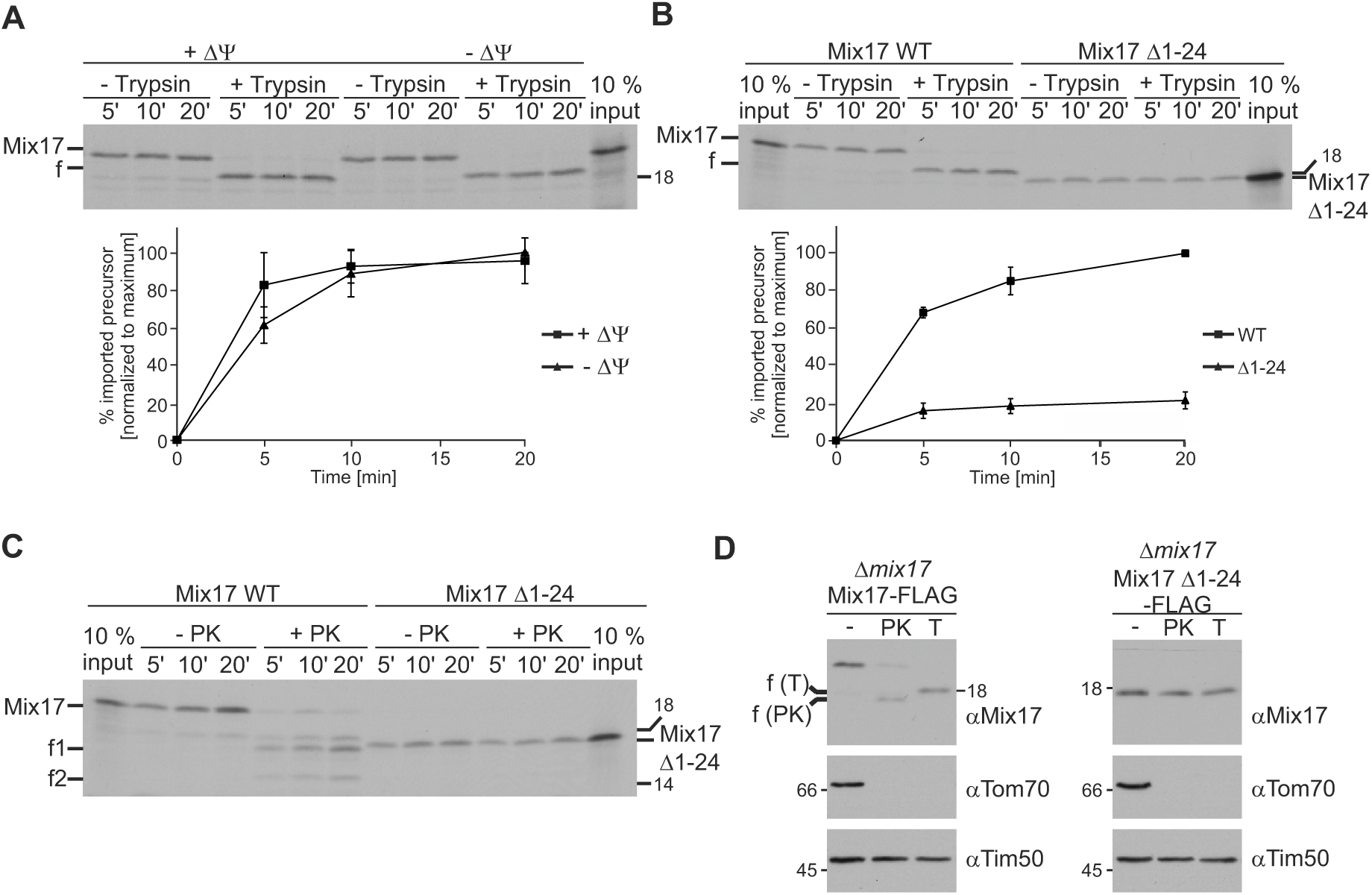
The N-terminus of Mix17 supports the import efficiency of Mix17 into mitochondria and is essential for its insertion into the outer membrane. **(A)** Import of Mix17 does not depend on mitochondrial membrane potential. ^35^S-labeled Mix17 protein was incubated with wild type mitochondria, which were either left untreated (+DY) or were treated with valinomycin (0.5 µM), antimycin A (8 µM) and oligomycin (20 µM) to dissipate the membrane potential (-DY). Samples were taken at the indicated time points and treated with trypsin as indicated. Mitochondria were re-isolated and the samples were analyzed by SDS-PAGE and autoradiography. (Upper panel) Autoradiograph of one representative experiment. (Lower panel) Quantitative analysis of 3 independent experiments. Results are presented as means of the imported protein normalized to the maximal imported material of -DY, which was set to 100%. Error bars, standard deviation. The full-length protein (Mix17) or the fragment generated by trypsin (f) are indicated **(B)** The N-terminal 24 residues of Mix17 strongly increase the import efficiency of Mix17. ^35^S-labeled full-length Mix17 and a mutant lacking the first 24 amino acids (Mix17 D1-24) were imported into wild type mitochondria. Samples were treated as in (A). (Upper panel) Autoradiograph of one representative experiment. (Lower panel) Quantitative analysis of 3 independently performed experiments. Results are presented as means of the imported protein in percent of the respective input and normalized to the maximal imported full-length Mix17, which was set to 100%. Error bars, standard deviation. **(C)** The N-terminus of Mix17 is essential for the insertion into the outer membrane *in vitro*. The import of full-length Mix17 and Mix17 D1-24 was performed as in (B) with the difference that PK was added, as indicated, instead of trypsin. The full-length protein (Mix17) or the fragments generated by PK (f1, f2) are indicated. The autoradiograph is representative of at least three repeats. **(D)** The N-terminus of Mix17 is essential for the insertion into the outer membrane *in vivo*. Mitochondria of yeast strains expressing full-length Mix17 WT-FLAG or Mix17 D1-24-FLAG were treated with PK or trypsin. Samples were analyzed by SDS-PAGE and immunoblotting using the indicated antibodies. The fragments generated by PK or trypsin are indicated. Blots are representative of 3 experiments.

In summary, we obtained evidence that the N-terminus of Mix17 is important for its integration into the outer membrane and supports the import of Mix17 into mitochondria.

### Mix17 can integrate into the outer membrane from the IMS

The ability of Mix17 Δ1-24 being imported into mitochondria but not exposed to the cytosol raises an intriguing question. Is full-length Mix17 first completely imported into mitochondria and then inserted into the outer membrane?

To address this question, we generated a Mix17 variant that is fused to the N-terminal bipartite presequence of the cytochrome *b*_2_ preprotein. This bipartite presequence drives import of the cyt*b*_2_ precursor protein by the TIM23 presequence translocase into mitochondria and the release of the precursor protein as intermediate form into the inner membrane (Hartl et al. 1987, Glick et al. 1992). The intermediate form is cleaved to release the mature form into the IMS (Schneider et al. 1991). Upon import of the radiolabeled Cyt*b*_2_(1-84)-Mix17 fusion protein into isolated mitochondria, we could detect an intermediate form and the release of Mix17 as mature form into the IMS. Importantly, the intermediate form was completely imported and protected from tryptic degradation, indicating that it is present in the IMS without exposing its N-terminus to the cytosol. The mature form, however, was entirely accessible to trypsin (Fig. 5), indicating its insertion into the outer membrane after the complete import of the fusion protein and the subsequent release of Mix17 into the IMS.

**Fig. 5.**
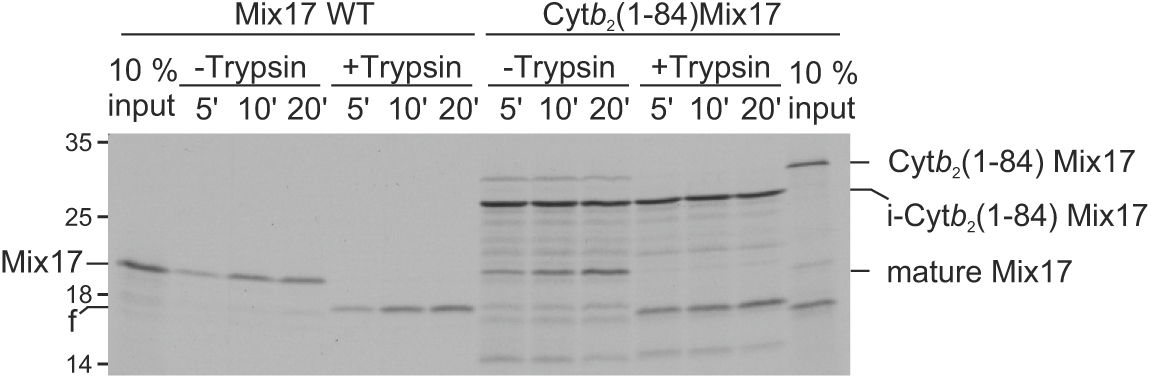
Mix17 is able to insert into the outer membrane from the intermembrane space. ^35^S-labeled full-length Mix17 or Mix17 fused to the N-terminus of Cyt*b*_2_ (Cyt*b*_2_(1-84) Mix17) were incubated with wild type mitochondria. The import was stopped at the indicated time points and the samples were treated with trypsin as indicated. Mitochondria were re-isolated and the samples were analyzed by SDS-PAGE and autoradiography. f, fragment generated by trypsin. i-Cyt*b*_2_(1-84) Mix17, intermediate form of Cyt*b*_2_(1-84) Mix17. The autoradiograph is representative of at least three repeats.

Thus, we conclude that Mix17 can insert into the outer membrane after being completely imported into the IMS.

### The insertion of Mix17 into the outer membrane does not depend on the TOB/SAM or MIM complexes

Two outer membrane complexes were shown to be able to insert proteins into the outer membrane, the TOB/SAM complex and the MIM complex (Kozjak et al. 2003, Paschen et al. 2003, Becker et al. 2008, Papic et al. 2011). Thus, we next analyzed whether one of these complexes might be responsible for the insertion of Mix17 into the outer membrane.

First, we tested a potential role of the TOB/SAM complex in the insertion process by depletion of its essential core component Tob55/Sam50. Virtually complete depletion of Tob55/Sam50 (Tob55↓ did not affect the steady state level of Mix17, although it already resulted in reduced level of the known substrates Tom40 and Porin (Fig. 6A). When we tested the protease accessibility of Mix17 in absence of Tob55, we also did not detect any difference compared to wild type (Fig. 6B). These results indicate that the TOB/SAM complex is not crucial for the membrane insertion of Mix17.

**Fig. 6.**
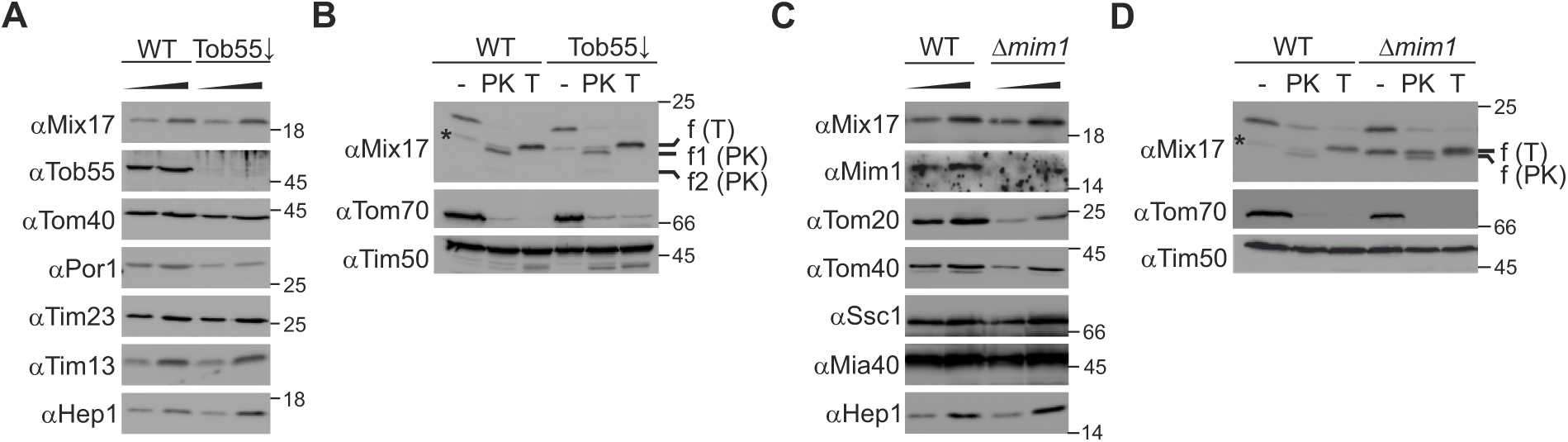
The insertion of Mix17 into the outer membrane does not depend on the TOB/SAM or MIM complexes. **(A)** The steady state level of Mix17 does not depend on the presence of Tob55. Mitochondria were isolated from wild type yeast and a mutant strain, in which Tob55 was downregulated (Tob55↓). Two different amounts of the mitochondria were analyzed by SDS-PAGE and immunoblotting using the indicated antibodies. **(B)** Tob55 is not important for the insertion of Mix17 into the outer membrane. Wild type and Tob55↓ mitochondria were either left untreated or incubated with PK or trypsin at isotonic conditions. Samples were analyzed by SDS-PAGE and immunoblotting. Star, degradation product of Mix17 in untreated mitochondria. The fragments generated by PK or trypsin are indicated. **(C)** The absence of Mim1 does not affect the level of Mix17. Mitochondria isolated from wild type yeast and a *MIM1* deletion mutant were analyzed as in (A). **(D)** Mim1 is dispensable for the insertion of Mix17 into the outer membrane. Wild type and D*mim1* mitochondria were analyzed as in (B). All blots are representative of three repeats.

Next, we analyzed whether the MIM complex is important for the topology of Mix17. Again, we did not detect reduced protein level of Mix17 in a *MIM1* deletion strain. The steady state level of Tom20, whose membrane integration is supported by Mim1, however, were strongly reduced (Fig. 6C). Moreover, Mix17 was accessible to proteases in mitochondria isolated from the *MIM1* deletion mutant similar to wild type mitochondria (Fig. 6D).

Based on these results, we conclude that neither the TOB/SAM complex nor the MIM complex play a crucial role in the insertion of Mix17 into the outer membrane.

### Mix17 interacts with Tom40

Our previous results suggested that Mix17 integrates into the outer membrane in a proteinaceous environment rather than into the lipid bilayer (Fig. 2A). Thus, we next asked whether the conserved hydrophobic segment of Mix17 is required for its membrane insertion. Therefore, we deleted the hydrophobic segment of Mix17 and tested whether this Mix17 Δ53-80 variant could span the outer membrane. Following its *in vitro* import, this variant was still accessible to PK and trypsin, indicating that the conserved hydrophobic segment is not required for the exposure of the N-terminus of Mix17 to the cytosol (Fig. 7A). In addition, this result supports our previous data, suggesting that Mix17 rather integrates into a protein complex than into the lipid bilayer of the outer membrane.

**Fig. 7.**
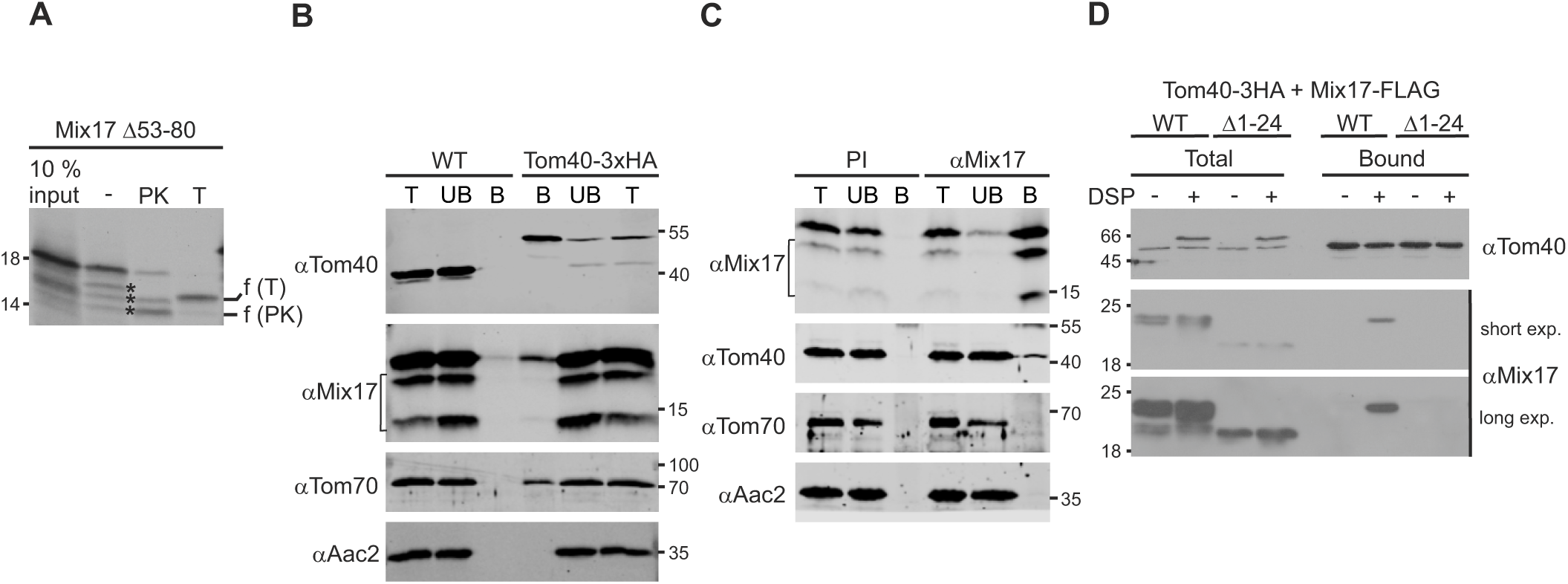
Mix17 interacts with Tom40 to be exposed to the cytosol. **(A)** The conserved hydrophobic stretch of Mix17 is not required for the insertion into the outer membrane. ^35^S-labeled Mix17 D53-80 protein was incubated with wild type mitochondria. Samples were either left untreated, or incubated with PK or trypsin (T). Re-isolated mitochondria were analyzed by SDS-PAGE and autoradiography. The fragments generated by PK or trypsin are indicated. Stars, potential degradation products of Mix17 in untreated sample. The autoradiograph is representative of at least three repeats. **(B, C)** Mix17 interacts with Tom40 at endogenous protein level. **(B)** DSP crosslinked mitochondria of wild type yeast and a yeast strain expressing Tom40-3xHA were lysed in digitonin containing buffer (1% w/v). Lysates were subjected to immunoprecipitation using anti-HAaffinity agarose. Samples were analyzed by SDS-PAGE and immunoblotting using the indicated antibodies. T, total lysate (4%); UB, unbound protein (4%); B, bound protein (100%). Bracket, degradation products of Mix17. **(C)** Wild type mitochondria were treated as described in (B) and subjected to immunoprecipitation using a Mix17 specific antibody or pre-immune serum bound to protein A sepharose. Samples were analyzed by SDS-PAGE and immunoblotting using the indicated antibodies. T, total lysate (4%); UB, unbound protein (4%); B, bound protein (100%). Bracket, degradation products of Mix17. **(D)** The N-terminus of Mix17 is essential for its interaction with Tom40. Mitochondria isolated from yeast strains expressing Tom40-3xHA and either full-length Mix17-FLAG or Mix17 D1-24-FLAG were left untreated or DSP crosslinked and subjected to immunoprecipitation as described in (B). Blots are representative of at least three repeats.

Since Mix17 did not integrate into the TOB/SAM or the MIM complex, Mix17 might use the Tom40 pore of the TOM complex to span the outer membrane. To address a role of Tom40 in the outer membrane insertion of Mix17, we asked whether Tom40 interacts with endogenous Mix17. To this end, we chromosomally tagged Tom40 with a C-terminal 3xHA-tag and performed an immunoprecipitation with anti-HA agarose beads. Strikingly, immunoprecipitation of Tom40-3xHA revealed the co-isolation of endogenous Mix17 from chemically crosslinked mitochondria. The co-isolation was specific, since Mix17 was not bound to the beads when wild type mitochondria were used. As control, the import receptor Tom70, a component of the TOM complex, was specifically co-purified with Tom40-3xHA, in contrast to Aac2, an inner membrane protein (Fig. 7B). To further substantiate the specificity of the interaction of Mix17 and Tom40, we applied immunoprecipitation of Mix17 (Fig. 7C). Interestingly, this again revealed the successful co-isolation of Tom40. Aac2 as well as Tom70, however, could not be co-purified, suggesting that Mix17 is most likely in close proximity to the protein import pore, but not to the receptor (Fig. 7C). Moreover, we found that the interaction could only be detected using chemically crosslinked mitochondria, indicating a transient rather than a stable interaction. We had observed that the N-terminal truncation variant Mix17 Δ1-24-FLAG is not protease accessible in mitochondria in contrast to Mix17-FLAG (Fig. 4D). Thus, we analyzed whether the N-terminal truncation variant interacts with Tom40. In consistency with the protease accessibility, we found that Mix17 Δ1-24-FLAG was not co-isolated with Tom40-3HA in contrast to the full-length protein (Fig. 7D). Of note, the steady state level of Mix17 Δ1-24-FLAG were considerably lower compared to the full-length variant, potentially due to the reduced import or stability of the N-terminal truncation mutant (Fig. 4B,C). In summary, Mix17 is able to interact with the TOM complex in the outer membrane at endogenous protein level. Mix17 Δ1-24-FLAG was not protease accessible in intact mitochondria and did not interact with Tom40, suggesting that the TOM complex mediates the insertion of the N-terminus of Mix17 into the outer membrane.

## Discussion

In the present study, we analyzed the submitochondrial location and the biogenesis of the highly conserved protein Mix17. We show that Mix17 spans the outer membrane with an N_out-_ C_in_-topology. Our data suggest a working model in which the N-terminus of Mix17 is important for the biogenesis of Mix17. First, it supports the import of Mix17 into the mitochondrial IMS (Fig. 8, step 1) where the disulfide bonds of Mix17 are formed by the MIA40 system (Fig. 8, step 2). Second, it is essential for the exposure to the cytosolic side of the mitochondrial outer membrane (Fig. 8, steps 3a,b). Mix17 interacts with Tom40, which is likely to be important for its exposure to the cytosol. This interaction might take place in the protein-conducting pore or at the protein-lipid interface of Tom40. Mix17 appears to associate with Tom40 rather dynamically than permanently (Fig. 8, steps 3a,b).

**Fig. 8.**
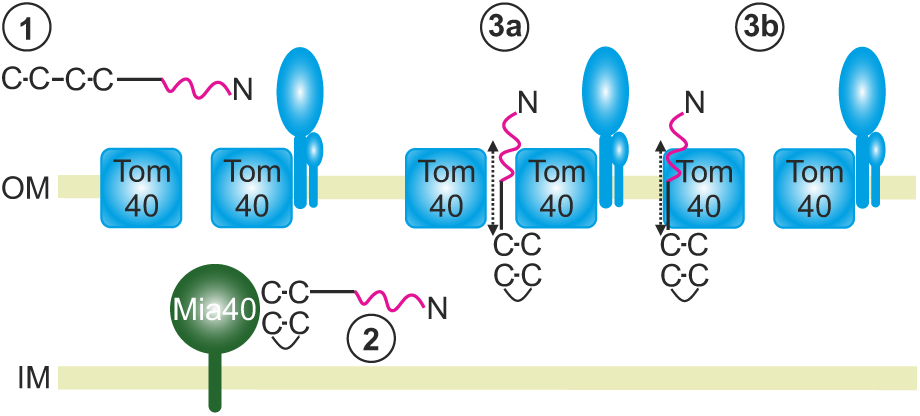
Working model of the Mix17 biogenesis. **(1)** The N-terminus of Mix17 supports the import of the protein into the mitochondrial IMS. **(2)** The MIA40 system is essential for the formation of the C-terminal disulfide bonds and thus the import of Mix17. **(3a,b)** The exposure of Mix17 to the cytosolic side of the mitochondrial outer membrane depends on its N-terminus, as wells as its association to Tom40. Two possible locations of Mix17 in proximity to Tom40 are shown. Dashed arrows indicate a potential dynamic topology of the Mix17 N-terminus. OM, outer membrane; IM, inner membrane.

Mix17 contains the twin Cx_9_C motif of a class of typical Mia40 substrates (Gabriel et al. 2007, Longen et al. 2009). We show that this motif is indeed crucial for import of Mix17. Moreover, we confirm previous reports of a Mia40-dependent import of Mix17 into mitochondria (Gabriel et al. 2007, Gornicka et al. 2014). However, we demonstrate here that Mix17 is the first Mia40 substrate that can insert into the outer membrane from the IMS to reach the cytosol. Of note, Mix17 has been previously described as a protease-resistant IMS protein (Gabriel et al. 2007, Gornicka et al. 2014). It is tempting to speculate that these different results are caused by individual experimental conditions or different states of mitochondria. A similar observation has been made before for Tim23, a core component of the TIM23 translocase. In contrast to previous studies (Kubrich et al. 1994), the N-terminus of Tim23 was reported to be protease accessible in intact mitochondria (Donzeau et al. 2000). It was suggested that this depends on the concentration of salt present in the experimental setup (Donzeau et al. 2000). In addition, the protease accessibility and, thus, the topology of Tim23 was shown to respond to the translocation activity of the TIM23 complex (Popov-Celeketic, 2008). Such a dependence of the topology on experimental and physiological conditions might also apply for Mix17.

Our results *in vitro* and *in vivo* demonstrate that the N-terminus of Mix17 is crucial for the insertion into the outer membrane. This appears to be independent of the presence of multiple positive charges in the N-terminus (data not shown). These positive charges are characteristic for mitochondrial targeting sequences. Likewise, Mix17 has been predicted to be a protein with such an amino-terminal presequence (Claros and Vincens 1996). However, in line with Gornicka et al. (2014) we observe that Mix17 did not require mitochondrial membrane potential for its import into mitochondria, in contrast to typical presequence proteins employing the TIM23 pathway. Moreover, the Mix17 C4S variant that could not use the Mia40 import pathway was not imported into mitochondria. Apparently, the N-terminus cannot function as a typical presequence sufficient to mediate import into mitochondria.

Nonetheless, we show that it strongly improves the import of Mix17, most likely due to its N-terminal positive charges. A similar observation was made for the import of the MICOS subunit Mic19. In contrast to typical Mia40 substrates, the presence of an unfolded DUF domain has been suggested to decrease the import efficiency of Mic19 (Ueda et al. 2019). In this case, myristoylation of the N-terminus counteracts this effect by mediating binding to the outer membrane and Tom20 (Ueda et al. 2019). Such inhibition of the import might also occur for Mix17 by the large segment amino-terminal to the CHCH domain. This segment is largely unfolded according to AlphaFold prediction (Jumper et al. 2021, Varadi et al. 2024). Interestingly, there are different results for CHCHD10/Mic14, the mammalian ortholog of Mix17, whether the N-terminus is required for import into mitochondria (Burstein et al. 2018, Lehmer et al. 2018). To address the function of mammalian Mic14, it will be crucial to elucidate whether CHCHD10/Mic14 also exposes its N-terminus across the outer membrane. Tom40 forms the central pore of the TOM complex, the main entry gate for import of proteins into mitochondria (Neupert and Herrmann 2007, Wiedemann and Pfanner 2017, Araiso et al. 2022). Interestingly, it has been reported that strongly overexpressed Mix17 can be co-purified with Tom40 during its import as a translocation intermediate (Gornicka et al. 2014). Our data suggest that endogenous Mix17 is embedded in the outer membrane in a proteinaceous environment after its import. Although we cannot rule out the existence of another so far unknown interaction partner of Mix17 in the mitochondrial outer membrane, we present first evidence that Tom40 might provide this proteinaceous environment to mediate the exposure of Mix17 to the cytosol. First, endogenous Mix17 interacts with Tom40. Second, the interaction with Tom40 was strongly reduced upon expression of the N-terminal truncation mutant Mix17 Δ1-24, which cannot be exposed to the cytosol. This interaction was only detected upon chemically crosslinking of mitochondria indicating a transient or dynamic rather than a stable interaction. In agreement with a dynamic interaction, a significant fraction of Mix17 was released from mitoplasts generated by swelling of mitochondria. Interestingly, Mix17 could be crosslinked to Tom40, but not to Tom70. These results are consistent with the hypothesis that Mix17 might span the TOM pore formed by Tom40. Still, it is possible that Mix17 interacts with Tom40 at the protein-lipid-interface of the TOM complex. Moreover, it cannot be excluded at this point that Tom40 might transiently mediate the insertion and assembly of Mix17 into its proteinaceous environment in the outer membrane. The mode of interaction of Mix17 with Tom40 and its effect on the import capacity of the TOM complex are important questions to be addressed in further studies.

Interestingly, the function of yeast Mix17 and its mammalian ortholog CHCHD10/Mic14 seems to be conserved since both are associated with respiration (Longen et al. 2009, Burstein et al. 2018, Lehmer et al. 2018). Thus, also the protein-protein interactions of Mix17 and CHCHD10/Mic14 might be conserved. In line with its enrichment at cristae junctions, CHCHD10/Mic14 was shown to interact with MICOS (Bannwarth et al. 2014, Genin et al. 2016). However, this interaction seems to be cell-type specific (Burstein et al. 2018). We show that Mix17 does not stably associate with MICOS but is in close proximity to Tom40. Interestingly, in yeast, MICOS was shown to interact with the TOM complex and to contribute to protein import (von der Malsburg et al. 2011, Bohnert et al. 2012, Zerbes et al. 2012). Therefore, it is tempting to speculate that Mix17 is also in the neighborhood of MICOS and might associate with the complex at specific conditions, perhaps to contribute to protein import. It will be an important task to analyze this and the factors contributing to its topology in future studies.

## Acknowledgements

M.E.H. was supported by *LMUexcellent* and the Deutsche Forschungsgemeinschaft (DFG), project number 413985647. K.H. was supported by the DFG project number 319071627. Furthermore, M.E.H. thanks Dr. Michael Kiebler, Ludwig-Maximilians-University, Munich, for generous and extensive support and Daniela Rieger for expert technical assistance and K.H. and M.R. thank Prof. Andreas Ladurner for his generous support and Petra Robisch and Alexandra Weinzierl for excellent technical assistance. Moreover, we thank Dejana Mokranjac for helpful discussion and reagents.

## Author contributions

M.R., J.S.F., K.D., and R.C., acquisition of data, analysis and interpretation of data, revising the article; K.H., conception and design, analysis and interpretation of data, drafting or revising the article; M.E.H., conception and design, acquisition of data, analysis and interpretation of data, drafting or revising the article.

## Data availability

All relevant data are included in the article and the Supplementary material. All strains, plasmids and antibodies used in this study can be obtained on request. Westerns Blots have been analyzed using Image Studio™ Lite software (LI-COR Biosciences; Version 5.2.5), Image Scanner III (GE) or ChemiDoc™ imaging system with Image Lab 6.1 software (Biorad).

## Material and Methods

### Yeast strains and cell growth

Chromosomal manipulations (knockouts, C-terminal tagging) were done as described before starting from YPH499 wild type (Longtine et al. 1998, Knop et al. 1999). The *MIX17* deletion strain was generated by replacing the entire coding region through marker cassette. In addition, the yeast wild type strain D273-10b was used. The genotypes of the strains used in this study are listed in Supplementary Table S1. All strains used in this study can be obtained on request.

For the generation of plasmids, the *MIX17* coding region was amplified by PCR. Point mutations were introduced by site-directed mutagenesis. Detailed information on the primers and restriction sites used are given in Supplementary Table 2S. Of note, for the *in vitro* synthesis of radiolabeled full-length Mix17, we exchanged at position 25 the amino acid methionine to alanine to omit the presence of an internal translation product that disturbs the analysis of the *in vitro* import assay. All plasmids used in this study can be obtained on request.

Yeast cells were grown on YP medium (1% w/v yeast extract, 2% peptone) supplemented with 3% glycerol (YPG), synthetic medium (0.67% yeast nitrogen base) supplemented with 2% glucose (SD), 3% galactose (SGal) or 3% glycerol (SG) or lactate medium (Daum et al. 1982, Izawa and Unger 2017). The yeast strain Mia40↓ carrying the *MIA40* gene under control of the *GAL10* promoter were first grown on galactose- and then transferred to glucose-containing medium for 16,5 h to deplete Mia40 from the cells. Tob55↓ cells were grown similarly, but shifted for 21 h to deplete Tob55. Yeast wild type cells and the respective mutants were grown in parallel and kept in logarithmic phase until further application.

### Isolation of mitochondria

Mitochondria were isolated according to established protocols (Daum et al. 1982, Izawa and Unger 2017). Cells were harvested by centrifugation and washed with distilled water. The cell pellet was resuspended in DTT containing buffer (10 mM final concentration) followed by an incubation for 10 min at 30°C with gentle agitation. Spheroplasts were generated by incubation with zymolyase (200 U per gram wet weight of cells) for 30 min at 30°C with gentle agitation. Sphaeroplasts were opened by pipetting or douncing in lysis buffer (20 mM MOPS-KOH pH 7.2 or 10mM Tris pH 7.4, and 1 mM EDTA, 0.6 M sorbitol, 0.2% (w/v) BSA, 1 mM phenylmethylsulfonyl fluoride (PMSF)). The crude mitochondrial fraction was harvested after a clarifying spin at 2,000 ×g and 4°C for 5 min by centrifugation at 14,000 ×g and 4°C for 10 min. Mitochondria were resuspended in isotonic SM buffer (0.6 M sorbitol, 20 mM MOPS, pH 7.4) or in SH buffer (0.6 M sorbitol, 20 mM HEPES, pH 7.4), frozen in liquid nitrogen and stored at −80°C.

### Alkaline extraction

Mitochondria were diluted in SM buffer to a concentration of 1 mg/ml. 100µg were mixed with 100µl 200 mM sodium carbonate and incubated for 30 min on ice. Soluble and membrane proteins were separated by centrifugation for 30 min at 91,000 ×g and 4°C. Soluble proteins in the supernatant were precipitated by addition of TCA (12%). The pelleted membrane proteins and the precipitated soluble proteins were resuspended in SDS sample buffer and analyzed by SDS-PAGE and immunoblotting (Fig. S2).

### Proteolytic accessibility assay

50 μg mitochondria were diluted in SM buffer (0.6 M sorbitol, 20 mM MOPS, pH 7.4), swelling buffer (20 mM MOPS, pH 7.4) or lysis buffer (1% (v/v) Triton X-100, 20 mM MOPS, pH 7.4). PK (final concentration: 12,5 µg/ml) or trypsin (final concentration: 12,5 or 25 µg/ml) were added as indicated and samples were incubated for 15 min on ice. Proteolysis was stopped by addition of PMSF (final concentration: 2 mM) or soybean trypsin inhibitor (final concentration: 500 µg/ml) followed by an incubation for 10 min on ice. Triton X-100 containing samples were precipitated with TCA. Mitochondria and mitoplasts were centrifuged at 17,000 ×g and 4°C for 20 min and the pellets were either first subjected to TCA precipitation and then analyzed or directly analyzed by SDS-PAGE and immunoblotting (Fig. S2).

### Import of radioactively labeled precursor protein

Radioactively labelled proteins were synthesized in the presence of [^35^S]methionine in a standard or in a transcription and translation coupled reticulocyte lysate system (Promega). Mitochondria (final concentration 0.5 mg/ml) were incubated in SI buffer (50 mM HEPES/KOH pH 7.2, 0.5 M sorbitol, 80 mM KCl, 10 mM MgAc, 2 mM KH_2_PO_4_, 2.5 mM EDTA, 1 mM MnCl_2_ and 0.01 % fatty acid-free BSA) containing 10 mM creatine phosphate, 0.1 mg/ml creatine kinase, 4 mM NADH and 2 mM ATP for 3 min at 12°C. Lysates were added and the reaction was further incubated at 12°C. At the indicated time points, samples were taken and diluted in ice-cold SH buffer (20 mM HEPES, 0,6 M sorbitol, pH 7.4) on ice. Mitochondria were pelleted and resuspended in SH buffer. Non-imported material was degraded by incubation with proteases (PK, 12,5µg/ml; trypsin, 25µg/ml) for 20 min on ice. Protease treatment was stopped by the addition of 2 mM PMSF or a 20-fold excess of soybean trypsin inhibitor, followed by an incubation for 5 min on ice. Mitochondria were re-isolated by centrifugation for 10 min at 14.000 ×g and 4°C and analyzed by SDS-PAGE and autoradiography (Fig. S2).

### Immunoprecipitation assay

When chemical crosslinking was performed prior to immunoprecipitation, 2 mg mitochondria were diluted in SI buffer (50 mM HEPES-KOH, 0.5 M sorbitol, 80 mM KCl, 10 mM Mg(Ac)_2_, 2 mM KH_2_PO_4_, 2.5 mM EDTA, 2.5 mM MnCl_2_, pH 7.2) to a concentration of 1 mg/ml. Dithiobissuccinimidylpropionate (DSP) was added (final concentration 800 μ followed by an incubation for 30 min on ice. The reaction was stopped by the addition of 1 M glycine pH 8.8 (final concentration of 160 mM) followed by an incubation for 10 min on ice.

Mitochondrial ATP was enzymatically depleted for 10 min at 25°C using 15 U hexokinase and 20 mM glucose or 10 U apyrase per 1 mg mitochondrial protein to protect Mix17 from unspecific degradation. Mitochondria were re-isolated and the pellets were lysed in IP buffer (50 mM Tris-HCl pH 7.4, 150 mM NaCl, 2 mM EDTA) containing 1% (w/v) digitonin and 2 mM PMSF. Cleared lysates were incubated with anti-HA agarose beads (30 μ beads/1 mg protein) or anti-Mix17 protein A sepharose beads (50 µl beads) for 1.5 h. The beads were washed three times with 500 µl IP buffer containing 0.1% (w/v) digitonin and 1 mM PMSF. Bound proteins were eluted with SDS sample buffer. Total lysate and unbound material were subjected to TCA precipitation to remove the digitonin, when proteins around 10 kDA should be detected. Otherwise, samples were directly diluted in SDS sample buffer. Samples were analyzed SDS-PAGE and immunoblotting (Fig. S2)

For immunoprecipitation of Mix17, anti-Mix17 antibody or pre-immune serum were bound to protein A sepharose beads over-night at 4°C. Beads were washed 3x with 0.5 ml 0.2 M boric acid (pH 9.0, NaOH). IgGs were coupled to protein A sepharose beads by incubation with 30 mM dimethylpimelimidate in 0.2M boric acid (pH 9.0, NaOH) for 30 min at room temperature. Beads were washed 3x with 0.5 ml 0.2 M ethanolamine (pH 8.0, HCl) and crosslinking was quenched by incubation in 1 ml 0.2 M ethanolamine (pH 8.0, HCl) for 2 h at room temperature. Afterwards beads were washed 2x with 1 ml IP buffer, 1x with 1ml 0.1 M glycine pH2.5, 2x with 0.5ml 0.1M Tris/HCl pH8.5, and 2x with 0.5 ml IP buffer.

### Antibody against Mix17

The antibody against Mix17 was generated by injecting a peptide comprising amino acids 99-118 into rabbits. For affinity purification of the Mix17 antibody, the antigen was coupled to SulfoLink Coupling Resin (1 mg protein/ml bead volume) in 50 mM Tris, 5 mM EDTA, pH 8.5. 6 ml antisera were diluted in 24 ml 10 mM Tris, pH 7.5 containing 1 mM PMSF, 1x complete protease inhibitor cocktail and 1 mM EDTA. The antisera were applied to the beads and washed successively with 10 mM Tris, pH 7.5 and 10 mM Tris, 0.5 M NaCl, pH 7.5. Elution of the antibody was performed in 100 mM sodium citrate, pH 4.0 followed by 100 mM glycine, pH 2.5. The individual fractions were immediately neutralized by addition of 1 M Tris, pH 8.8. The specificity of the antibody was tested using samples generated from the *MIX17* deletion strain or strains expressing tagged versions of Mix17 and can be seen in Fig. 4D. All additional antibodies used in this study are listed in Supplementary Table S3.

## References

Alkhaja, A. K., D. C. Jans, M. Nikolov, M. Vukotic, O. Lytovchenko, F. Ludewig, W. Schliebs, D. Riedel, H. Urlaub, S. Jakobs and M. Deckers (2012). MINOS1 is a conserved component of mitofilin complexes and required for mitochondrial function and cristae organization. Mol Biol Cell 23, 247–257. 10.1091/mbc.E11-09-0774.

Araiso, Y., K. Imai and T. Endo (2022). Role of the TOM Complex in Protein Import into Mitochondria: Structural Views. Annu Rev Biochem 91, 679–703. 10.1146/annurev-biochem-032620-104527.

Bannwarth, S., S. Ait-El-Mkadem, A. Chaussenot, E. C. Genin, S. Lacas-Gervais, K. Fragaki, L. Berg-Alonso, Y. Kageyama, V. Serre, D. G. Moore, A. Verschueren, C. Rouzier, I. Le Ber, G. Auge, C. Cochaud, F. Lespinasse, K. N’Guyen, A. de Septenville, A. Brice, P. Yu-Wai-Man, H. Sesaki, J. Pouget and V. Paquis-Flucklinger (2014). A mitochondrial origin for frontotemporal dementia and amyotrophic lateral sclerosis through CHCHD10 involvement. Brain 137, 2329–2345. 10.1093/brain/awu138.

Becker, T., S. Pfannschmidt, B. Guiard, D. Stojanovski, D. Milenkovic, S. Kutik, N. Pfanner, C. Meisinger and N. Wiedemann (2008). Biogenesis of the mitochondrial TOM complex: Mim1 promotes insertion and assembly of signal-anchored receptors. J Biol Chem 283, 120–127. 10.1074/jbc.M706997200.

Burstein, S. R., F. Valsecchi, H. Kawamata, M. Bourens, R. Zeng, A. Zuberi, T. A. Milner, S. M. Cloonan, C. Lutz, A. Barrientos and G. Manfredi (2018). In vitro and in vivo studies of the ALS-FTLD protein CHCHD10 reveal novel mitochondrial topology and protein interactions. Hum Mol Genet 27, 160–177. 10.1093/hmg/ddx397.

Claros, M. G. and P. Vincens (1996). Computational method to predict mitochondrially imported proteins and their targeting sequences. Eur J Biochem 241, 779–786. 10.1111/j.1432-1033.1996.00779.x.

Daum, G., S. M. Gasser and G. Schatz (1982). Import of proteins into mitochondria. Energy-dependent, two-step processing of the intermembrane space enzyme cytochrome b2 by isolated yeast mitochondria. J. Biol. Chem. 257, 13075–13080.

Donzeau, M., K. Kaldi, A. Adam, S. Paschen, G. Wanner, B. Guiard, M. F. Bauer, W. Neupert and M. Brunner (2000). Tim23 links the inner and outer mitochondrial membranes. Cell 101, 401–412.

Edwards, R., S. Gerlich and K. Tokatlidis (2020). The biogenesis of mitochondrial intermembrane space proteins. Biol Chem 401, 737–747. 10.1515/hsz-2020-0114.

Gabriel, K., D. Milenkovic, A. Chacinska, J. Muller, B. Guiard, N. Pfanner and C. Meisinger (2007). Novel mitochondrial intermembrane space proteins as substrates of the MIA import pathway. J Mol Biol 365, 612–620. 10.1016/j.jmb.2006.10.038.

Genin, E. C., M. Plutino, S. Bannwarth, E. Villa, E. Cisneros-Barroso, M. Roy, B. Ortega-Vila, K. Fragaki, F. Lespinasse, E. Pinero-Martos, G. Auge, D. Moore, F. Burte, S. Lacas-Gervais, Y. Kageyama, K. Itoh, P. Yu-Wai-Man, H. Sesaki, J. E. Ricci, C. Vives-Bauza and V. Paquis-Flucklinger (2016). CHCHD10 mutations promote loss of mitochondrial cristae junctions with impaired mitochondrial genome maintenance and inhibition of apoptosis. EMBO Mol Med 8, 58–72. 10.15252/emmm.201505496.

Glick, B. S., A. Brandt, K. Cunningham, S. Muller, R. L. Hallberg and G. Schatz (1992). Cytochromes c1 and b2 are sorted to the intermembrane space of yeast mitochondria by a stop-transfer mechanism. Cell 69, 809–822.

Gornicka, A., P. Bragoszewski, P. Chroscicki, L. S. Wenz, C. Schulz, P. Rehling and A. Chacinska (2014). A discrete pathway for the transfer of intermembrane space proteins across the outer membrane of mitochondria. Mol Biol Cell 25, 3999–4009. 10.1091/mbc.E14-06-1155.

Harner, M., C. Korner, D. Walther, D. Mokranjac, J. Kaesmacher, U. Welsch, J. Griffith, M. Mann, F. Reggiori and W. Neupert (2011). The mitochondrial contact site complex, a determinant of mitochondrial architecture. EMBO J 30, 4356–4370. 10.1038/emboj.2011.379.

Hartl, F. U., J. Ostermann, B. Guiard and W. Neupert (1987). Successive translocation into and out of the mitochondrial matrix: targeting of proteins to the intermembrane space by a bipartite signal peptide. Cell 51, 1027–1037.

Hell, K. (2008). The Erv1-Mia40 disulfide relay system in the intermembrane space of mitochondria. Biochim Biophys Acta 1783, 601–609. 10.1016/j.bbamcr.2007.12.005.

Hoppins, S., S. R. Collins, A. Cassidy-Stone, E. Hummel, R. M. Devay, L. L. Lackner, B. Westermann, M. Schuldiner, J. S. Weissman and J. Nunnari (2011). A mitochondrial-focused genetic interaction map reveals a scaffold-like complex required for inner membrane organization in mitochondria. J Cell Biol 195, 323–340. 10.1083/jcb.201107053.

Izawa, T. and A. K. Unger (2017). Isolation of Mitochondria from Saccharomyces cerevisiae. Methods Mol Biol 1567, 33–42. 10.1007/978-1-4939-6824-4_3.

John, G. B., Y. Shang, L. Li, C. Renken, C. A. Mannella, J. M. Selker, L. Rangell, M. J. Bennett and J. Zha (2005). The mitochondrial inner membrane protein mitofilin controls cristae morphology. Mol Biol Cell 16, 1543–1554. 10.1091/mbc.E04-08-0697.

Jumper, J., R. Evans, A. Pritzel, T. Green, M. Figurnov, O. Ronneberger, K. Tunyasuvunakool, R. Bates, A. Zidek, A. Potapenko, A. Bridgland, C. Meyer, S. A. A. Kohl, A. J. Ballard, A. Cowie, B. Romera-Paredes, S. Nikolov, R. Jain, J. Adler, T. Back, S. Petersen, D. Reiman, E. Clancy, M. Zielinski, M. Steinegger, M. Pacholska, T. Berghammer, S. Bodenstein, D. Silver, O. Vinyals, A. W. Senior, K. Kavukcuoglu, P. Kohli and D. Hassabis (2021). Highly accurate protein structure prediction with AlphaFold. Nature 596, 583–589. 10.1038/s41586-021-03819-2.

Knop, M., K. Siegers, G. Pereira, W. Zachariae, B. Winsor, K. Nasmyth and E. Schiebel (1999). Epitope tagging of yeast genes using a PCR-based strategy: more tags and improved practical routines. Yeast 15, 963–972. 10.1002/(SICI)1097-0061(199907)15:10B<963::AID-YEA399>3.0.CO;2-W.

Koehler, C. M. and H. L. Tienson (2009). Redox regulation of protein folding in the mitochondrial intermembrane space. Biochim Biophys Acta 1793, 139–145. 10.1016/j.bbamcr.2008.08.002.

Kozjak, V., N. Wiedemann, D. Milenkovic, C. Lohaus, H. E. Meyer, B. Guiard, C. Meisinger and N. Pfanner (2003). An essential role of Sam50 in the protein sorting and assembly machinery of the mitochondrial outer membrane. J Biol Chem 278, 48520–48523. 10.1074/jbc.C300442200.

Kubrich, M., P. Keil, J. Rassow, P. J. Dekker, J. Blom, M. Meijer and N. Pfanner (1994). The polytopic mitochondrial inner membrane proteins MIM17 and MIM23 operate at the same preprotein import site. FEBS Lett. 349, 222–228.

Lehmer, C., M. H. Schludi, L. Ransom, J. Greiling, M. Junghanel, N. Exner, H. Riemenschneider, J. van der Zee, C. Van Broeckhoven, P. Weydt, M. T. Heneka and D. Edbauer (2018). A novel CHCHD10 mutation implicates a Mia40-dependent mitochondrial import deficit in ALS. EMBO Mol Med 10. 10.15252/emmm.201708558.

Longen, S., M. Bien, K. Bihlmaier, C. Kloeppel, F. Kauff, M. Hammermeister, B. Westermann, J. M. Herrmann and J. Riemer (2009). Systematic analysis of the twin cx(9)c protein family. J Mol Biol 393, 356–368. 10.1016/j.jmb.2009.08.041.

Longtine, M. S., A. McKenzie, 3rd, D. J. Demarini, N. G. Shah, A. Wach, A. Brachat, P. Philippsen and J. R. Pringle (1998). Additional modules for versatile and economical PCR-based gene deletion and modification in Saccharomyces cerevisiae. Yeast 14, 953–961.

Mesecke, N., N. Terziyska, C. Kozany, F. Baumann, W. Neupert, K. Hell and J. M. Herrmann (2005). A disulfide relay system in the intermembrane space of mitochondria that mediates protein import. Cell 121, 1059–1069. 10.1016/j.cell.2005.04.011.

Neupert, W. and J. M. Herrmann (2007). Translocation of proteins into mitochondria. Annu Rev Biochem 76, 723–749. 10.1146/annurev.biochem.76.052705.163409.

Papic, D., K. Krumpe, J. Dukanovic, K. S. Dimmer and D. Rapaport (2011). Multispan mitochondrial outer membrane protein Ugo1 follows a unique Mim1-dependent import pathway. J Cell Biol 194, 397–405. 10.1083/jcb.201102041.

Paschen, S. A., T. Waizenegger, T. Stan, M. Preuss, M. Cyrklaff, K. Hell, D. Rapaport and W. Neupert (2003). Evolutionary conservation of biogenesis of beta-barrel membrane proteins. Nature 426, 862–866. 10.1038/nature02208.

Rabl, R., V. Soubannier, R. Scholz, F. Vogel, N. Mendl, A. Vasiljev-Neumeyer, C. Korner, R. Jagasia, T. Keil, W. Baumeister, M. Cyrklaff, W. Neupert and A. S. Reichert (2009). Formation of cristae and crista junctions in mitochondria depends on antagonism between Fcj1 and Su e/g. J Cell Biol 185, 1047–1063. 10.1083/jcb.200811099.

Riemer, J., N. Bulleid and J. M. Herrmann (2009). Disulfide formation in the ER and mitochondria: two solutions to a common process. Science 324, 1284–1287. 10.1126/science.1170653.

Schneider, A., M. Behrens, P. Scherer, E. Pratje, G. Michaelis and G. Schatz (1991). Inner membrane protease I, an enzyme mediating intramitochondrial protein sorting in yeast. EMBO J. 10, 247–254.

Stojanovski, D., P. Bragoszewski and A. Chacinska (2012). The MIA pathway: a tight bond between protein transport and oxidative folding in mitochondria. Biochim Biophys Acta 1823, 1142–1150. 10.1016/j.bbamcr.2012.04.014.

Ueda, E., Y. Tamura, H. Sakaue, S. Kawano, C. Kakuta, S. Matsumoto and T. Endo (2019). Myristoyl group-aided protein import into the mitochondrial intermembrane space. Sci Rep 9, 1185. 10.1038/s41598-018-38016-1.

Varadi, M., D. Bertoni, P. Magana, U. Paramval, I. Pidruchna, M. Radhakrishnan, M. Tsenkov, S. Nair, M. Mirdita, J. Yeo, O. Kovalevskiy, K. Tunyasuvunakool, A. Laydon, A. Zidek, H. Tomlinson, D. Hariharan, J. Abrahamson, T. Green, J. Jumper, E. Birney, M. Steinegger, D. Hassabis and S. Velankar (2024). AlphaFold Protein Structure Database in 2024: providing structure coverage for over 214 million protein sequences. Nucleic Acids Res 52, D368–D375. 10.1093/nar/gkad1011.

von der Malsburg, K., J. M. Muller, M. Bohnert, S. Oeljeklaus, P. Kwiatkowska, T. Becker, A. Loniewska-Lwowska, S. Wiese, S. Rao, D. Milenkovic, D. P. Hutu, R. M. Zerbes, A. Schulze-Specking, H. E. Meyer, J. C. Martinou, S. Rospert, P. Rehling, C. Meisinger, M. Veenhuis, B. Warscheid, I. J. van der Klei, N. Pfanner, A. Chacinska and M. van der Laan (2011). Dual role of mitofilin in mitochondrial membrane organization and protein biogenesis. Dev Cell 21, 694–707. 10.1016/j.devcel.2011.08.026.

Wiedemann, N. and N. Pfanner (2017). Mitochondrial Machineries for Protein Import and Assembly. Annu Rev Biochem 86, 685–714. 10.1146/annurev-biochem-060815-014352.

