## Supplementary Information for "The Mia40 substrate Mix17 exposes its N-terminus to the cytosolic side of the mitochondrial outer membrane"

```
H.sapiens         -----------------MPR--GSR-SAASRPASRPAAPSAHPPAHPPPSAAAPAPAPSG 40
R.norvegicus      -----------------MPR--GSR-SAAARPASRPA----HPPAHPPPSAPAPAPATSG 36
C.elegans         MVRRRTASPSPSAPVRSAPRPAAQSSFAAPPPRPAAAAPAYHPPAAPTP-MGAPMGAPSQ 59
N.crassa          MPRQSRGSARPSVPARK--------PVAPTNQQQQRPASTYAPPAAAPHAPPAAAAPVSQ 52
S.cerevisiae      -MARSRGSSRPISRSRPTQTRSASTMAAPVHPQQQQQPNAYS--------HPPAAGAQTR 51
                                             *                              :

H.sapiens         QPGLMAQMATTAAGVAVGSAVGHVMGSALTGAFSGGSSEPSQP-----AVQQAPTPAAPQ 95
R.norvegicus      QPGLMAQMASTAAGVAVGSAVGHVMGSALTSAFSGGSSEPAQP-----AVQQAPARPASH 91
C.elegans         GPGLMKQMAATAGGVAIGSAVGHAVGGMFTGGGSSH--AEQAP-----AAAAAPAGAPQA 112
N.crassa          GPGLFGQMASTAAGVAIGSSIGHAI----GGMFSGGGSSAAPEAAAAPVQ--AQAAAAQN 106
S.cerevisiae      QPGMFAQMASTAAGVAVGSTIGHTLGAGITGMFSGSGSDSAPVEQQQQNMANTSGQTQTD 111
                   **:: ***:**.***:**::**.:     .  *.                 :

H.sapiens         PLQMGPCAYEIRQFLDCST-TQSDLSLCEGFSEALKQCKYYHGLSSLP 142
R.norvegicus      PLQMGPCAYEIKQFLDCST-TQSDLTLCEGFSEALKQCKYNHGLSSLP 138
C.elegans         SGYSQPCEFEWRQFVDCAQ-NQSDVSLCNGFNDIFKQCKARYA----- 154
N.crassa          SSWGNNCSEATKSFTQCMDQHQGNMQICGWYLEQLKACQAAASQY--- 151
S.cerevisiae      QQLGRTCEIDARNFTRCLDENNGNFQICDYYLQQLKACQEAARQY*-- 156
                        *    :.*  *    :.:. :*  : : :* *:
```

**Fig. S1. Mix17 is conserved throughout evolution**

The amino acid sequence of *Saccharomyces cerevisiae* Mix17 and its orthologs in *Homo sapiens*, *Rattus norvegicus*, *Caenorhabditis elegans*, and *Neurospora crassa* were aligned using Clustal Omega (Madeira et al. 2022). Bolt, conserved hydrophobic segment; underlined, immunogenic peptide used for antibody generation; box, conserved $CX_9C$ motif.

**Supplementary Table 1.** ***S. cerevisiae*** **strains used in this study.**

| Strain name | Genotype | Reference |
|---|---|---|
| YPH499 | *MAT*a *ade2-101 his3-Δ200 leu2- trp1-Δ63 ura3-52 lys2-801* | (Sikorski and Hieter 1989) |
| D273-10B | *MAT*α *mal GAL* | (Sherman 1963) |
| Mix17-3xMyc | YPH499 *MIX17::3xMyc::HIS3MX6* | This study |
| Tom40-3xHA | YPH499 *TOM40-3xHA::HIS3MX6* | This study |
| Mic10-3xHA | YPH499 *MIC10-3xHA::HIS3MX6* | (Harner et al. 2011) |
| Mic60-3xHA | YPH499 *MIC60-3xHA::HIS3MX6* | (Harner et al. 2011) |
| Δ*mix17* | YPH499 *mix17Δ::LEU2 (K. lactis)* | This study |
| Δ*mim1* | YPH499 *mim1Δ::LEU2 (K. lactis)* | This study |
| Mia40↓ | YPH499 *kanMX4::*$P_{GAL10}$ *MIA40* | (Terziyska et al. 2005) |
| Tob55↓ | YPH499 *kanMX4::*$P_{GAL10}$ *TOB55* | (Paschen et al. 2003) |

**Supplementary Table 2. Vector constructs and primers used in this study.**

| Construct | Primer name | Restriction site | Sequence |
|---|---|---|---|
| pYX233 Mix17-FLAG | RI ACC Mix17 | EcoRI | ctcgaattcaccatggcacgttcaagaggatcatc |
| | Mix17 XhoI nSt | XhoI | ctcctcgaggtattgacgtgcagcttcctggcaggc |
| pYX233 Mix17 Δ1-24-FLAG | RI ACC Mix17 Δ24 | EcoRI | ctcgaattcaccatggcggctccagttca |
| | Mix17 XhoI nSt | XhoI | ctcctcgaggtattgacgtgcagcttcctggcaggc |
| pGEM4 Mix17C4S | Mix17 CS 1 | - | aaactttacacgttctttggatgaaaacaacggc |
| | Mix17 CS 2 | - | ctcgcgtctatttcagaagttcttcccaactgctg |
| | Mix17 CS 3 | - | caactaaaagcctcccaggaagctgcacgtcaa |
| | Mix17 CS 4 | - | tgcaagtaataatcagatatctggaagttgccgttg |
| pGEM4 Mix17 | pGEM4 Mix17_for | Xma1 | gaatacacggaattcgagctcggtacccgggatggcacgttcaagagg |
| | pGEM4 Mix17_rev | Pst1 | tatagggagaccggaagcttgcatgcctgcagttagtattgacgtgcagc |
| | Mix17 M25A for | - | cgcttctaccgcggcggctccag |
| | Mix17 M25A rev | - | gaccttgtctgagtaggtc |
| pGEM4 Mix17 Δ1-24 | pGEM4 Mix17Δ24_for | Xma1 | gaatacacggaattcgagctcggtacccgggtccgcttctaccatggc |
| | pGEM4 Mix17_rev | Pst1 | tatagggagaccggaagcttgcatgcctgcagttagtattgacgtgcagc |
| pGEM4 Mix17 Δ53-80 | Mix17 Δ53-80 f | - | accggtatgttttccggatc |
| | Mix17 Δ53-80 r | - | ctgtctcgtttgggcacc |
| pGEM4 Cyt$b_2$(1-84) Mix17 | Mix17 XhoI | XhoI | ctcctcgagttagtattgacgtgcagcttcctg |
| | BglII Mix17 | BglII | ctcagatctatggcacgttcaagaggatcatc |
| | KpnI Cyt$b_2$ | KpnI | ctcggtacccgggatgctaaaatacaaacctttactaaaaatc |
| | Mix17 HindIII | HindIII | gaccggaagcttgcatgcctgcagttagtattgacgtgcagc |

**Supplementary Table 3. Antibodies used in this study.**

| Antibody | Source | Identifier/Reference | Dilution |
|---|---|---|---|
| IRDye 680RD Goat anti-Mouse IgG | LI-COR | Cat. #962-68070 | 1:10,000 |
| IRDye 800CW Goat anti-Rabbit IgG | LI-COR | Cat. #926-32211 | 1:10,000 |
| Goat Anti-Rabbit IgG-HRP Conjugate | BIO-RAD | Cat. #170-6515 | 1:10,000 |
| Mouse monoclonal anti-Myc | Roche | Cat. #ROAMYC | 1:250 |
| Rabbit polyclonal anti-Mix17 | LMU Munich | This study | 1:250 |
| Mouse monoclonal anti-HA | Santa Cruz | Cat. #sc-57592 | 1:250 |
| Rabbit polyclonal anti-Tim13 | LMU Munich | (Paschen et al. 2000) | 1:200 |
| Rabbit polyclonal anti-Tim23 | LMU Munich | (Mokranjac et al. 2003) | 1:500 |
| Rabbit polyclonal anti-Tim50 | LMU Munich | (Mokranjac et al. 2003) | 1:500 |
| Rabbit polyclonal anti-Tom20 | LMU Munich | (Krimmer et al. 2001) | 1:500 |
| Rabbit polyclonal anti-Tom40 | LMU Munich | (Kiebler et al. 1990) | 1:1,000 |
| Rabbit polyclonal anti-Tom70 | LMU Munich | (Schlossmann et al. 1996) | 1:500 |
| Rabbit polyclonal anti-Tob55 | LMU Munich | (Paschen et al. 2003) | 1:250 |
| Rabbit polyclonal anti-Ssc1 | LMU Munich | (Sichting et al. 2005) | 1:1000 |
| Rabbit polyclonal anti- Mia40 | LMU Munich | (Terziyska et al. 2005) | 1:500 |
| Rabbit polyclonal anti- Mic10 | LMU Munich | (Harner et al. 2011) | 1:250 |
| Rabbit polyclonal anti- Mic26 | LMU Munich | (Harner et al. 2011) | 1:250 |
| Rabbit polyclonal anti- Mic27 | LMU Munich | (Harner et al. 2011) | 1:250 |
| Rabbit polyclonal anti- Mic60 | LMU Munich | (Rabl et al. 2009) | 1:250 |
| Rabbit polyclonal anti-Mim1 | LMU Munich | (Waizenegger et al. 2005) | 1:250 |
| Rabbit polyclonal anti-Hep1 | LMU Munich | (Sichting et al. 2005) | 1:250 |
| Rabbit polyclonal anti-Por1 | LMU Munich | (Kleene et al. 1987) | 1:1,000 |
| Rabbit polyclonal anti-Aco1 | LMU Munich | (Adam et al. 2006) | 1:1,000 |
| Rabbit polyclonal anti-Aac2 | LMU Munich | (Sollner et al. 1990) | 1:1,000 |

### Supplementary References


Adam, A. C., C. Bornhovd, H. Prokisch, W. Neupert and K. Hell (2006). The Nfs1 interacting protein Isd11 has an essential role in Fe/S cluster biogenesis in mitochondria. EMBO J *25*, 174-183. 10.1038/sj.emboj.7600905.

Harner, M., C. Korner, D. Walther, D. Mokranjac, J. Kaesmacher, U. Welsch, J. Griffith, M. Mann, F. Reggiori and W. Neupert (2011). The mitochondrial contact site complex, a determinant of mitochondrial architecture. EMBO J *30*, 4356-4370. 10.1038/emboj.2011.379.

Kiebler, M., R. Pfaller, T. Sollner, G. Griffiths, H. Horstmann, N. Pfanner and W. Neupert (1990). Identification of a mitochondrial receptor complex required for recognition and membrane insertion of precursor proteins. Nature *348*, 610-616.

Kleene, R., N. Pfanner, R. Pfaller, T. A. Link, W. Sebald, W. Neupert and M. Tropschug (1987). Mitochondrial porin of Neurospora crassa: cDNA cloning, in vitro expression and import into mitochondria. EMBO J *6*, 2627-2633. 10.1002/j.1460-2075.1987.tb02553.x.

Krimmer, T., D. Rapaport, M. T. Ryan, C. Meisinger, C. K. Kassenbrock, E. Blachly-Dyson, M. Forte, M. G. Douglas, W. Neupert, F. E. Nargang and N. Pfanner (2001). Biogenesis of porin of the outer mitochondrial membrane involves an import pathway via receptors and the general import pore of the TOM complex. J Cell Biol *152*, 289-300. 10.1083/jcb.152.2.289.

Madeira, F., M. Pearce, A. R. N. Tivey, P. Basutkar, J. Lee, O. Edbali, N. Madhusoodanan, A. Kolesnikov and R. Lopez (2022). Search and sequence analysis tools services from EMBL-EBI in 2022. Nucleic Acids Res *50*, W276-W279. 10.1093/nar/gkac240.

Mokranjac, D., S. A. Paschen, C. Kozany, H. Prokisch, S. C. Hoppins, F. E. Nargang, W. Neupert and K. Hell (2003). Tim50, a novel component of the TIM23 preprotein translocase of mitochondria. EMBO J. *22*, 816-825. 10.1093/emboj/cdg090.

Paschen, S. A., U. Rothbauer, K. Kaldi, M. F. Bauer, W. Neupert and M. Brunner (2000). The role of the TIM8-13 complex in the import of Tim23 into mitochondria. EMBO J *19*, 6392-6400. 10.1093/emboj/19.23.6392.

Paschen, S. A., T. Waizenegger, T. Stan, M. Preuss, M. Cyrklaff, K. Hell, D. Rapaport and W. Neupert (2003). Evolutionary conservation of biogenesis of beta-barrel membrane proteins. Nature *426*, 862-866. 10.1038/nature02208.
Rabl, R., V. Soubannier, R. Scholz, F. Vogel, N. Mendl, A. Vasiljev-Neumeyer, C. Korner, R. Jagasia, T. Keil, W. Baumeister, M. Cyrklaff, W. Neupert and A. S. Reichert (2009). Formation of cristae and crista junctions in mitochondria depends on antagonism between Fcj1 and Su e/g. J Cell Biol *185*, 1047-1063. 10.1083/jcb.200811099.
Schlossmann, J., R. Lill, W. Neupert and D. A. Court (1996). Tom71, a novel homologue of the mitochondrial preprotein receptor Tom70. J. Biol. Chem. *271*, 17890-17895. 10.1074/jbc.271.30.17890.
Sherman, F. (1963). Respiration-deficient mutants of yeast. I. Genetics. Genetics *48*, 375-385. 10.1093/genetics/48.3.375.
Sichting, M., D. Mokranjac, A. Azem, W. Neupert and K. Hell (2005). Maintenance of structure and function of mitochondrial Hsp70 chaperones requires the chaperone Hep1. EMBO J *24*, 1046-1056. 10.1038/sj.emboj.7600580.
Sikorski, R. S. and P. Hieter (1989). A system of shuttle vectors and yeast host strains designed for efficient manipulation of DNA in Saccharomyces cerevisiae. Genetics *122*, 19-27. 10.1093/genetics/122.1.19.
Sollner, T., R. Pfaller, G. Griffiths, N. Pfanner and W. Neupert (1990). A mitochondrial import receptor for the ADP/ATP carrier. Cell *62*, 107-115. 0092-8674(90)90244-9 [pii].
Terziyska, N., T. Lutz, C. Kozany, D. Mokranjac, N. Mesecke, W. Neupert, J. M. Herrmann and K. Hell (2005). Mia40, a novel factor for protein import into the intermembrane space of mitochondria is able to bind metal ions. FEBS Lett *579*, 179-184. 10.1016/j.febslet.2004.11.072.
Waizenegger, T., S. Schmitt, J. Zivkovic, W. Neupert and D. Rapaport (2005). Mim1, a protein required for the assembly of the TOM complex of mitochondria. EMBO Rep *6*, 57-62. 10.1038/sj.embor.7400318.
